# A transcriptome-wide Mendelian randomization study to uncover tissue-dependent regulatory mechanisms across the human phenome

**DOI:** 10.1101/563379

**Authors:** Tom G Richardson, Gibran Hemani, Tom R Gaunt, Caroline L Relton, George Davey Smith

## Abstract

**Background:** Developing insight into tissue-specific transcriptional mechanisms can help improve our understanding of how genetic variants exert their effects on complex traits and disease. By applying the principles of Mendelian randomization, we have undertaken a systematic analysis to evaluate transcriptome-wide associations between gene expression across 48 different tissue types and 395 complex traits.

**Results:** Overall, we identified 100,025 gene-trait associations based on conventional genome-wide corrections (P < 5 × 10^−08^) that also provided evidence of genetic colocalization. These results indicated that genetic variants which influence gene expression levels in multiple tissues are more likely to influence multiple complex traits. We identified many examples of tissue-specific effects, such as genetically-predicted *TPO*, *NR3C2* and *SPATA13* expression only associating with thyroid disease in thyroid tissue. Additionally, *FBN2* expression was associated with both cardiovascular and lung function traits, but only when analysed in heart and lung tissue respectively.

We also demonstrate that conducting phenome-wide evaluations of our results can help flag adverse on-target side effects for therapeutic intervention, as well as propose drug repositioning opportunities. Moreover, we find that exploring the tissue-dependency of associations identified by genome-wide association studies (GWAS) can help elucidate the causal genes and tissues responsible for effects, as well as uncover putative novel associations.

**Conclusions:** The atlas of tissue-dependent associations we have constructed should prove extremely valuable to future studies investigating the genetic determinants of complex disease. The follow-up analyses we have performed in this study are merely a guide for future research. Conducting similar evaluations can be undertaken systematically at http://mrcieu.mrsoftware.org/Tissue_MR_atlas/.

## Introduction

Advancements in high-throughput sequencing technologies present an unprecedented opportunity to investigate the molecular determinants of complex disease. This has facilitated the identification of genetic variants that influence gene expression, known as expression quantitative trait loci (eQTL). Recent studies have demonstrated the benefit of using eQTL data to help understand the underlying mechanisms of findings from genome-wide association studies (GWAS)^1–3^. Moreover, endeavours leveraging eQTL data derived from different tissue types can help to further ascertain the biological and clinical relevance of variants associated with complex traits^4–6^. In particular, these efforts are important when investigating tissue specificity, the phenomenon whereby a gene’s function is restricted to particular tissue types^7^.

An important challenge in molecular epidemiology is assessing how associations between gene expression and complex traits depend upon the tissue analysed. We previously proposed an analytical pipeline to detect associations between tissue-specific gene expression and complex traits by applying the principles of Mendelian randomization (MR)^8–10^. This approach harnesses eQTL as instrumental variables to investigate whether genetic variants at a locus influence both gene expression and complex trait variation. Furthermore, this framework has advantages over alternative transcriptome-wide approaches by incorporating techniques of genetic colocalization^11, 12^. This helps to mitigate the likelihood of spurious findings attributed to two separate but correlated variants at a locus, one responsible for influencing gene expression and the other affecting the associated complex trait. As such, associations supported by evidence of genetic colocalization are more likely to be driven by a shared genetic factor. Crucially, we note that genetic colocalization is necessary, but not sufficient, for causality. This is because the genetic effect may influence the associated trait due to mediated changes in gene expression, or it may operate on both through independent biological pathways^13^.

In this study, we have applied our framework to comprehensively evaluate the association between the transcription of 32,116 protein-coding, RNA- and pseudo-genes and 395 complex traits. This was undertaken across 48 tissue types using data from the GTEx consortium^14^ (v7), as well as whole blood derived data from the eQTLGen project^15^ (n=31,684). With this putative causal map of tissue-dependent associations we have undertaken several extensive analyses. Firstly, we have evaluated the relationship between gene expression across many tissues and pleiotropy; the phenomenon whereby a gene influences variation in multiple traits^16^. Next, we undertook a series of transcriptome and phenome-wide analyses to uncover tissue-dependent associations. Findings such as these can help to develop insight into the underlying regulatory mechanisms which reside along the causal pathway from a genetic variant to its associated complex trait. Moreover, they can help uncover pleiotropic effects that may be confined to separate tissue types.

We also demonstrate that phenome-wide evaluations of target genes have translatable value. For example, they can help predict whether therapeutic intervention will result in potential on-target side effects, as well as propose novel scope for drug repurposing. This is particularly attractive given previous evidence has reported that genetic associations supporting therapeutic intervention can improve efficacy and safety rates^17, 18^. Finally, we have explored the tissue-dependency of associations between selected genetic variants detected by GWAS for blood pressure traits. Our findings suggest that integrating tissue-specific eQTL data can help prioritise likely functional genes and tissues responsible for GWAS signals.

## Results

### Constructing an atlas of tissue-dependent associations across the human phenome

We pooled together eQTL data from the GTEx consortium (v7) for 48 tissue types (n=80 to 491, Supplementary Table 1) and the eQTLGen project using findings derived from whole blood (n=31,684). Full summary statistics for 395 complex traits were obtained from large-scale GWAS (Supplementary Table 2). To investigate the association between the transcription of up to 32,116 genes (i.e. protein-coding, RNA- and pseudo-genes) and each trait in turn, we applied two-sample summary Mendelian randomization (2SMR)^19^ and assessed genetic colocalization using the heterogeneity in dependent instruments (HEIDI) method (v0.710)^2^. A lenient p-value threshold of P < 1.0 × 10^−04^ was used to define lead eQTL as instrumental variables in our analysis. However, this threshold is simply a heuristic for highlighting associations worthy of follow-up^20^. Investigations of results can therefore apply a more (or less) stringent threshold by filtering associations based on the p-value for lead eQTL in analyses. All findings can be visualised and downloaded using our web application located at http://mrcieu.mrsoftware.org/Tissue_MR_atlas/. A schematic of our study analysis can be found in Figure 1.

**Figure 1.**
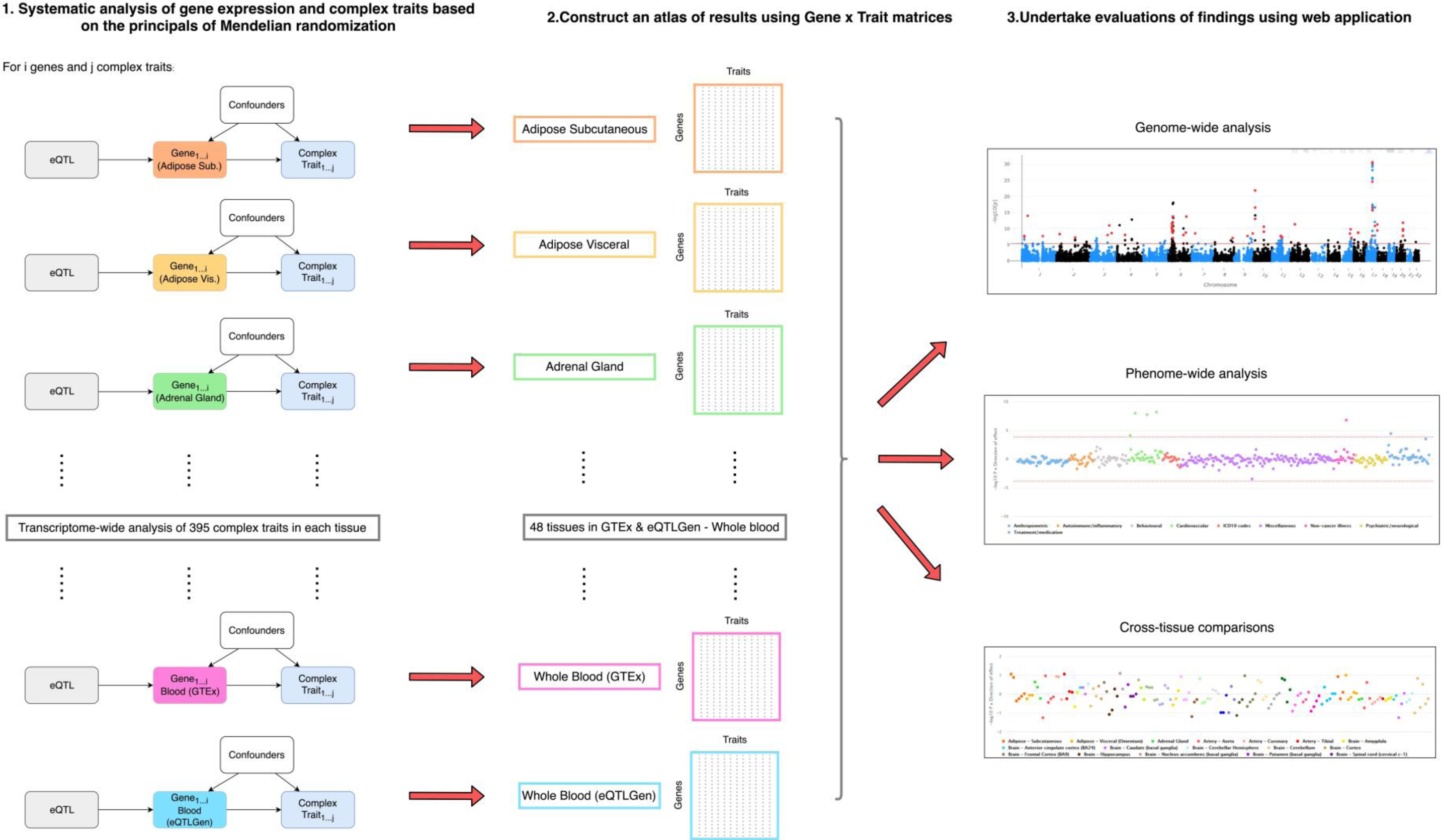
A schematic of the analysis plan in this study

Each analysis undertaken was adjusted for conventional genome-wide corrections (i.e. MR P < 5.0 × 10^−08^) and filtered for evidence of genetic colocalization (i.e. HEIDI P > 0.05/number of associations detected). In total, 100,025 MR associations were robust to multiple testing and genetic colocalization based on these criteria. We also found that associations derived using eQTLGen data were strongly enriched for associations using GTEx whole blood data compared to different tissue types from this resource (P<1.0×10^−04^).

We hypothesised that variants which influence gene expression levels in multiple tissues are more likely to influence multiple complex traits. To investigate this, we firstly grouped associations according to the organ that tissues were derived from (Supplementary Table 3). The reason for this is because we may expect similar association signals to be shared between tissues in GTEx which were part of the same embryonic tissue during development. For example, the various types of brain tissue from the GTEx consortium (e.g. Amygdala, Cerebellum etc.) were allocated to the ‘Brain’ tissue group. This was to reduce false positive findings from effectively counting the same association twice (e.g. gene expression in various types of brain tissue associated with the same neurological trait).

We identified strong evidence of a positive relationship between the number of associated traits for each lead eQTL and the number of tissues they were detected in (Beta=0.60, SE=0.02, P<1.0×10^−16^). This analysis was adjusted for minor allele frequencies, linkage disequilibrium (LD) score and distance to gene expression probe for lead eQTL, given that these genomic properties may influence the number of associated traits for a given SNP. In a subsequent analysis we clustered eQTL effects based of their associated genes. Overall, there was a positive correlation between the number of traits that each gene was associated with and the number of different tissue groups that these associations were detected across (r^2^= 0.38, Figure 2).

**Figure 2.**
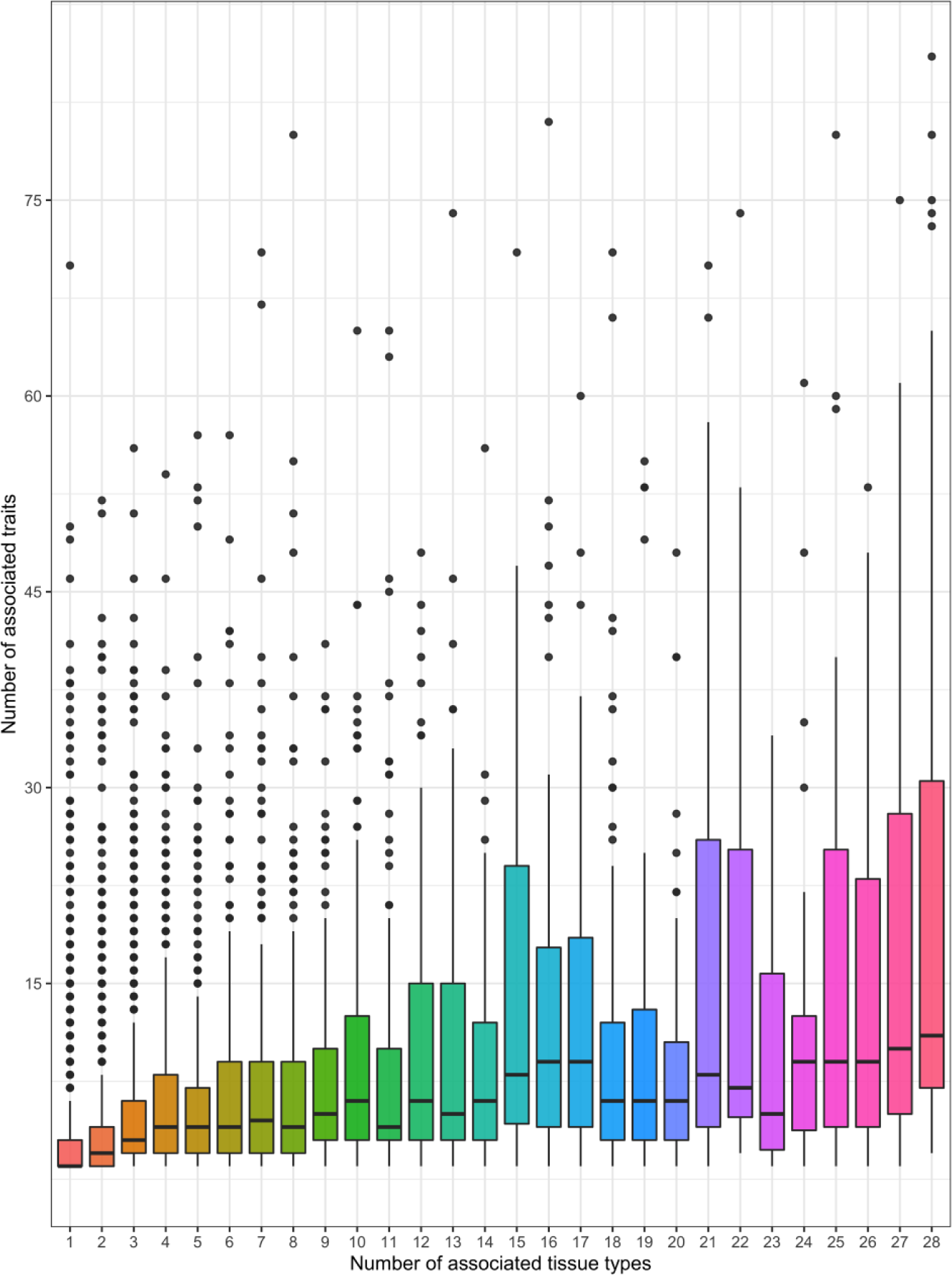
Box plot portraying the correlation in our atlas that genetically determined gene expression is more likely to be associated with multiple traits when expressed across multiple diverse tissue types.

### A transcriptome-wide evaluation of thyroid disease to uncover tissue-dependent effects

Findings from our extensive analyses can be used to conduct hypothesis-driven investigations of tissue-dependent effects. For example, we hypothesised that genetic variants which influence risk of thyroid disease (defined as self-reported hypothyroidism or myxoedema in the UK Biobank study) may likely act via changes to gene expression in thyroid tissue. Figure 3 illustrates the results of a transcriptome-wide evaluation between thyroid-derived gene expression and thyroid disease using results from our atlas. We identified 58 associations which survived multiple testing (P < 5.66 × 10^−06^, i.e. 0.05/8834 test) and 33 of these survived HEIDI filtering (P > 8.62 × 10^−04^ based on 0.05/58 tests) (Supplementary Table 4). However, 12 of these were in the HLA region and should be interpreted with caution due to the extensive linkage disequilibrium which may hinder the reliability of genetic colocalization analyses^21^.

**Figure 3.**
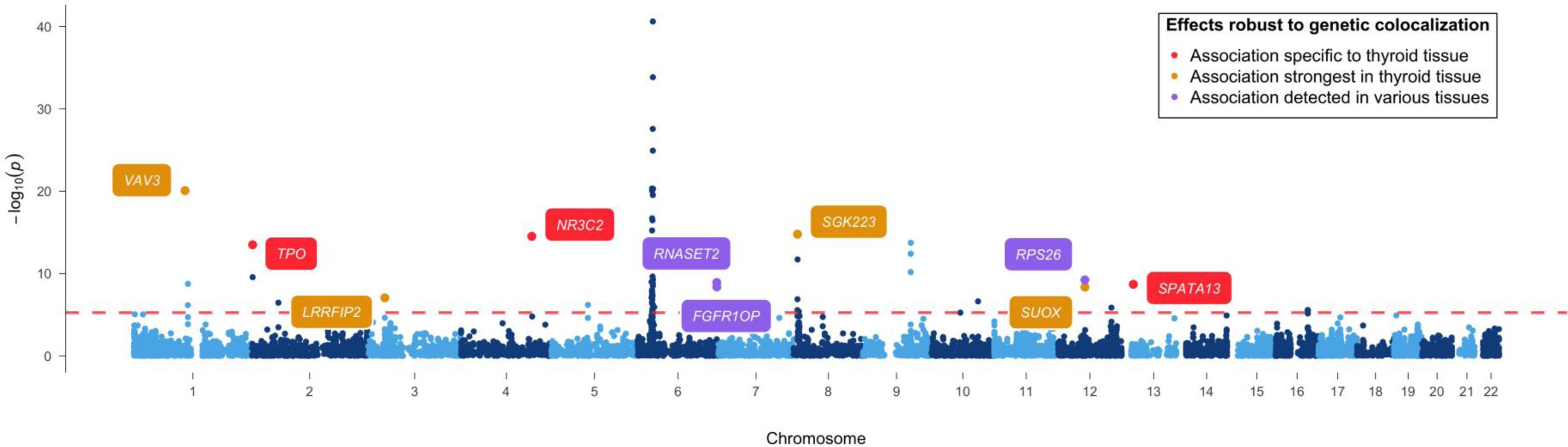
A Manhattan plot illustrating the association between genetically influenced gene expression derived from thyroid tissue and self-reported thyroid disease in the UK Biobank study. Amongst signals which were robust to genetic colocalization we identified associations only detected using thyroid tissue (red), associations detected with the strongest evidence in thyroid tissue (i.e. evidence of association in at least 2 tissues with thyroid being the strongest – yellow) and associations observed across many different tissue types (i.e. evidence of association in at least 2 tissues where thyroid is not the strongest – purple).

We evaluated the association for each of these genetic effects on thyroid disease in all other available tissue types. Although we report these genetic effects based on their corresponding gene symbols, it should be noted that they are based on the MR effect estimates using lead eQTL. We found that in particular 3 of these associations appeared to be highly tissue-specific (*TPO*, *NR3C2* and *SPATA13*) as they were only identified in thyroid tissue after correcting for the number of tissues evaluated (Supplementary Tables 5–7). Cross-tissue associations for *TPO* and thyroid disease are illustrated in Figure 4a. These effects provided strong evidence of heterogeneity (Cochran’s Q statistic=104.8, P=7.12×10^−14^), which reflects the tissue-dependency of associations for *TPO*.

**Figure 4.**
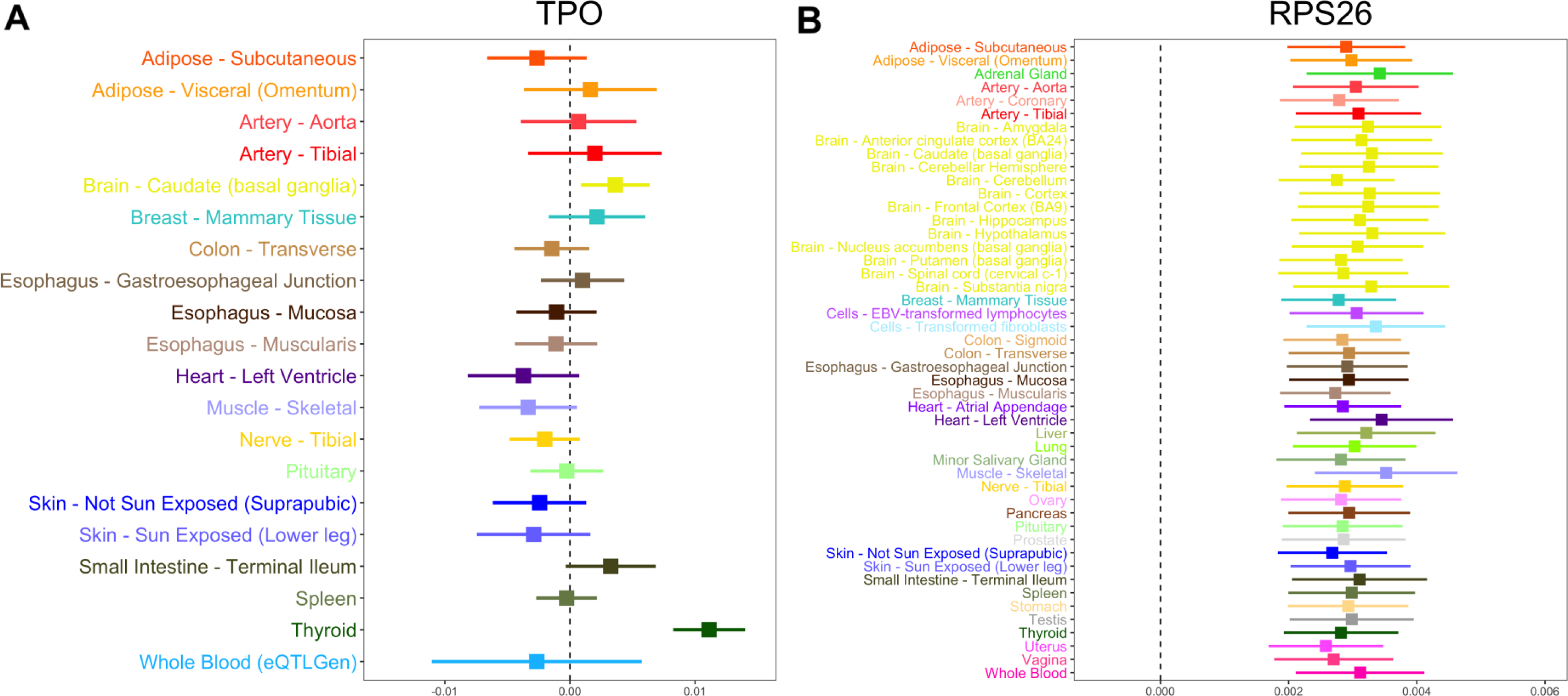
Forest plots for tissue-dependent effects identified in our analysis between genes associated with thyroid disease. a) The association for *TPO* appeared to be tissue-dependent and most strongly associated in thyroid tissue, b) whereas *RPS26* expression was strongly associated in all tissues assessed. The horizontal lines in these plots indicates the null of beta=0.

We also identified effects detected most strongly in thyroid tissue, although evidence of association was still identified in other tissue types (*VAV3*, *LRRFIP2*, *SGK223* and *SUOX*, Supplementary Table 8–11). These results also demonstrate that certain associations appear to be detected across many or all tissue types assessed. For example, the association between *RPS26* and thyroid disease was detected across all 48 tissue types assessed as portrayed in Figure 4b (Supplementary Table 12). In contrast to *TPO*, there was weak evidence of heterogeneity for *RPS26* (Cochran’s Q statistic=27.1, P=0.99), reflecting consistent associations across all tissues analysed.

### Conducting phenome-wide association analyses to evaluate tissue-dependent effects

Along with evaluating our results in a transcriptome-wide manner as above, exploring findings in a phenome-wide manner can be a powerful approach to explore pleiotropy. As a demonstration of this, in the previous analysis we ascertained that *RPS26* is strongly associated with thyroid disease across many different tissues. Undertaking a phenome-wide scan of this gene’s expression using whole blood suggests that the corresponding variant used as an instrument is highly pleiotropic, as a total of 48 associations survived multiple testing and HEIDI corrections (Supplementary Table 13 and Figure 5a). *RPS26* therefore appears to be a case in point that genes expressed in many tissues may be more likely to influence multiple different phenotypes.

**Figure 5.**
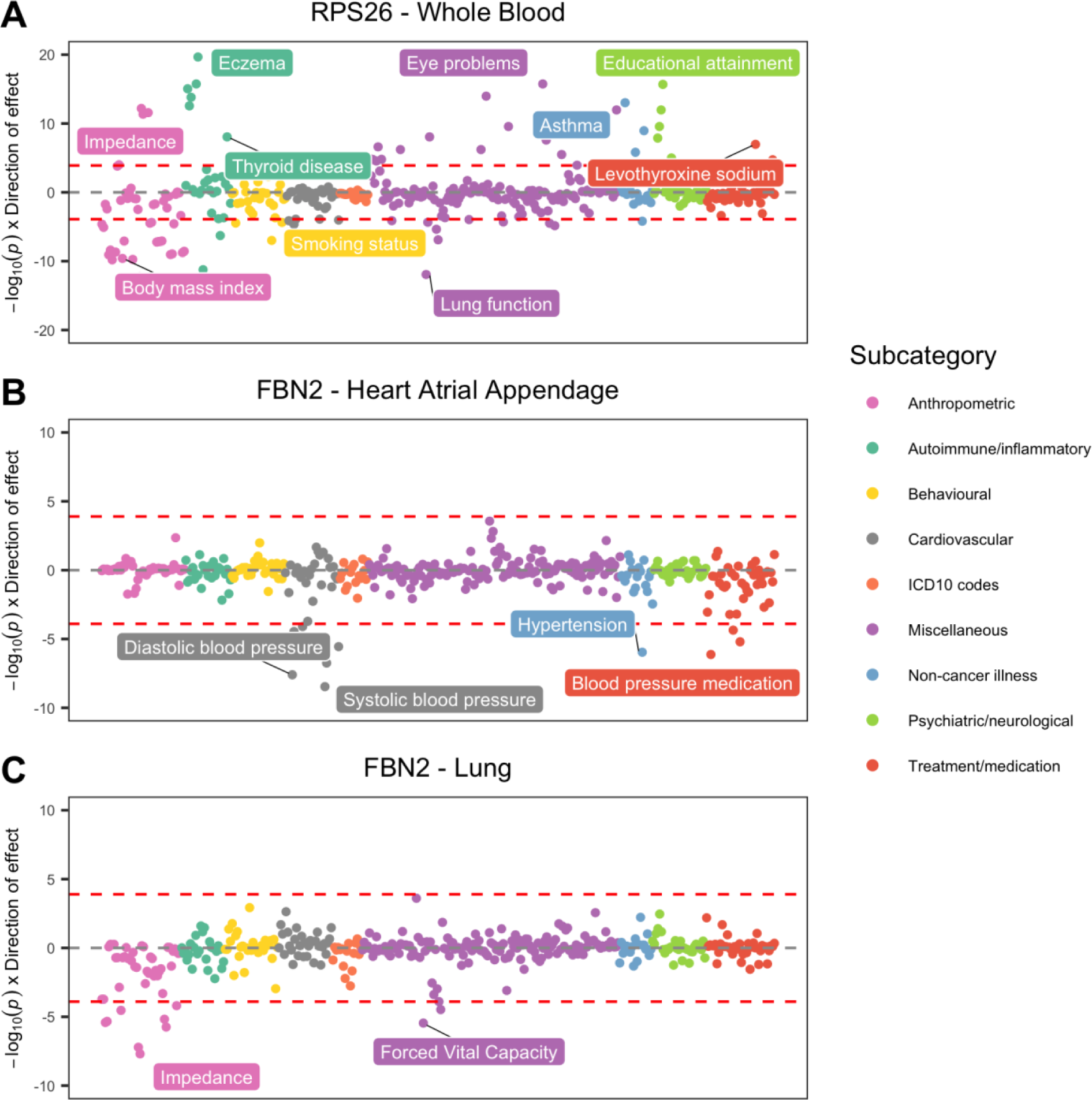
Miami plots illustrating phenome-wide associations between genes in different tissue types. a) *RPS26* expression derived from whole blood was associated with many diverse traits, b) *FBN2* expression derived from heart tissue was associated with blood pressure traits, c) *FBN2* associations with blood pressure attenuated when analysed using lung-derived data. However, other associations (e.g. measures of lung function) were observed instead.

Investigating phenome-wide associations for genes of interest can also yield insight into tissue-dependent effects. As an example, we evaluated genes in our atlas associated with two traits with a substantial heritable component within the UK Biobank study; diastolic blood pressure and forced vital capacity (FVC). We found that *FBN2* expression was linked with both traits in our results, although when using heart tissue derived data only the effects on blood pressure were observed (Supplementary Table 14 and Figure 5b). However, these associations attenuate when investigating this effect in other tissues types. Moreover, when evaluating phenome-wide associations of *FBN2* using lung tissue-derived eQTL data we identified evidence of association with FVC (MR P = 3.51 × 10^−06^, Supplementary Table 15 and Figure 5c). Findings such as this may be attributed to different eQTL used as instrumental variables for the same gene but within a different tissue type (as is the case for *FBN2*). As such, they may elucidate tissue-dependent regulatory mechanisms that can help explain associations at pleiotropic loci^22^.

### Harnessing findings to highlight potential side effects of therapeutic intervention and drug repositioning opportunities

Exploring our associations in a phenome-wide manner may also be valuable for other purposes, such as helping validate whether genes may be viable drug targets^23^. A well-established example of this is the impact of HMG-coenzyme A reductase (HMG-CoA) inhibition using statins, which is known to reduce low-density lipoprotein (LDL) cholesterol levels. However, this is known to also potentially result in increased bodyweight and risk of diabetes^24^.

Undertaking a phenome-wide evaluation of *HMGCR* (the gene responsible for HMG-CoA) using data derived from skeletal muscle tissue supports these findings. After removing associations which did not survive HEIDI corrections, we observed strong positive associations between the lead eQTL for this gene and high LDL and total cholesterol levels (Supplementary Table 16 and Figure 6a). We also identified evidence of association with lower body mass index (MR P = 1.63 × 10^−05^), although the association with self-reported diabetes did not survive phenome-wide corrections (MR P = 0.002). Nonetheless, these findings help support the notion that Mendelian randomization analyses can help mimic the findings of randomized control trials^25^ and identify potential on-target side effects of therapeutic intervention^26^. We note however that the tissue analysed may play an important part in such analyses, particularly with respect to the sensitivity of genetic colocalization. For instance, repeating evaluations of *HMGCR* using whole blood data derived from eQTLGen suggests that associations signals are less robust to colocalization in this tissue (e.g. HEIDI P = 1.8×10^−06^ with LDL cholesterol). In general however, cross-tissue comparisons of our results need to be interpreted with caution due to the differing sample sizes of eQTL datasets derived from the GTEx consortium.

**Figure 6.**
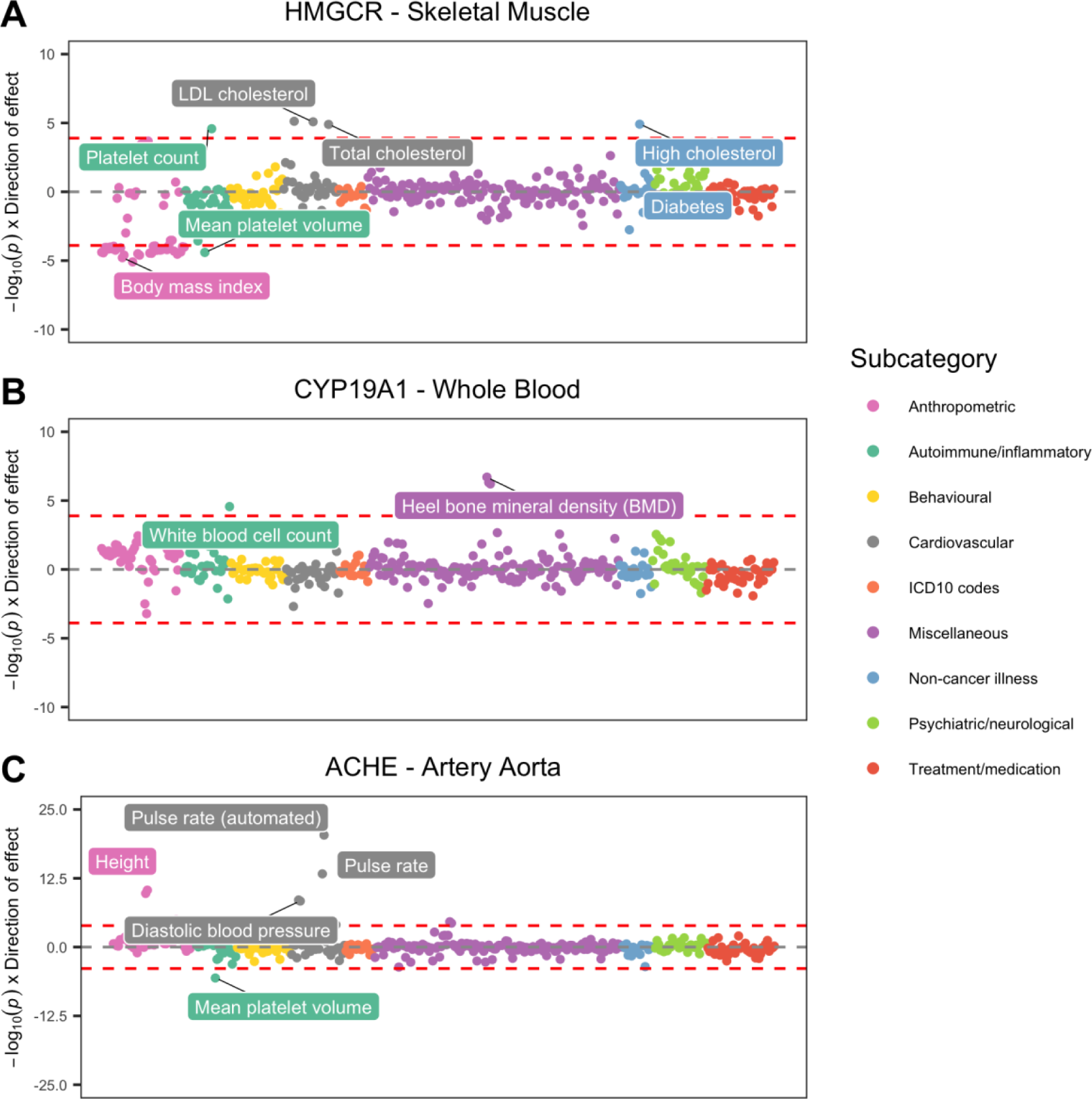
Miami plots representing phenome-wide associations between genes targeted for therapeutic intervention. a) *HMGCR* associations reflect known consequences of statins, b) *CYP19A1* associations support adverse on-target side effects on bone mineral density, c) *ACHE* associations demonstrate scope for novel repurposing opportunities (e.g. possible inhibition to reduce blood pressure).

A more novel demonstration of highlighting potential adverse effects was identified by conducting a similar analysis for *CYP19A1* expression using data derived from whole blood (Supplementary Table 17 and Figure 6b). This gene has been previously targeted using the drug Anastrozole to reduce risk of breast cancer^27^, although reported side effects include increased risk of osteoporosis^28^. Our phenome-wide scan of *CYP19A1* provided evidence of this reported on-target adverse effect, as we identified strong evidence of association with heel bone mineral density (BMD) (MR P = 1.96 × 10^−07^).

Conducting these types of evaluations may also be beneficial for potential drug repositioning opportunities. For instance, *ACHE*, which is a target for drugs used to treat cognitive decline in Alzheimer’s patients, such as Galantamine and Donepezil^29^. The causal pathway targeted by these drugs would likely be expected to inhibit *ACHE* expression in brain tissue. However, conducting a phenome-wide evaluation for this gene in other tissues (such as artery aorta) indicates that its transcription is associated with higher blood pressure (Supplementary Table 18 and Figure 6c). Further research could therefore explore whether inhibiting this gene’s product may have beneficial implications for hypertension.

### Leveraging tissue-specific expression data to help elucidate genes responsible for association signals

An important challenge in genetic epidemiology is pinpointing the causal gene responsible for association signals detected by GWAS. This is a complex problem for several reasons, including the co-expression that can exist between nearby genes that is often difficult to disentangle^30^. We previously proposed that integrating tissue-specific eQTL data with findings from GWAS may help with such endeavours^9^.

For example, rs7500448 is strongly associated with diastolic blood pressure (DBP) (after adjustment for medication) based on analyses undertaken using data from the UK Biobank study (P=6.3×10^−15^). Harnessing all available tissue-dependent results from our atlas allowed us to evaluate associations between nearby genes for which this SNP is an eQTL. Doing so identified only one association signal which survived multiple comparisons, which was *CDH13* using eQTL data derived from the aorta (MR P=2.78×10^−08^) (Supplementary Table 19 and Figure 7a). This provides strong evidence that *CDH13* may be the causal gene responsible for this effect, and that its expression in the aorta may play a role in blood pressure variation.

**Figure 7.**
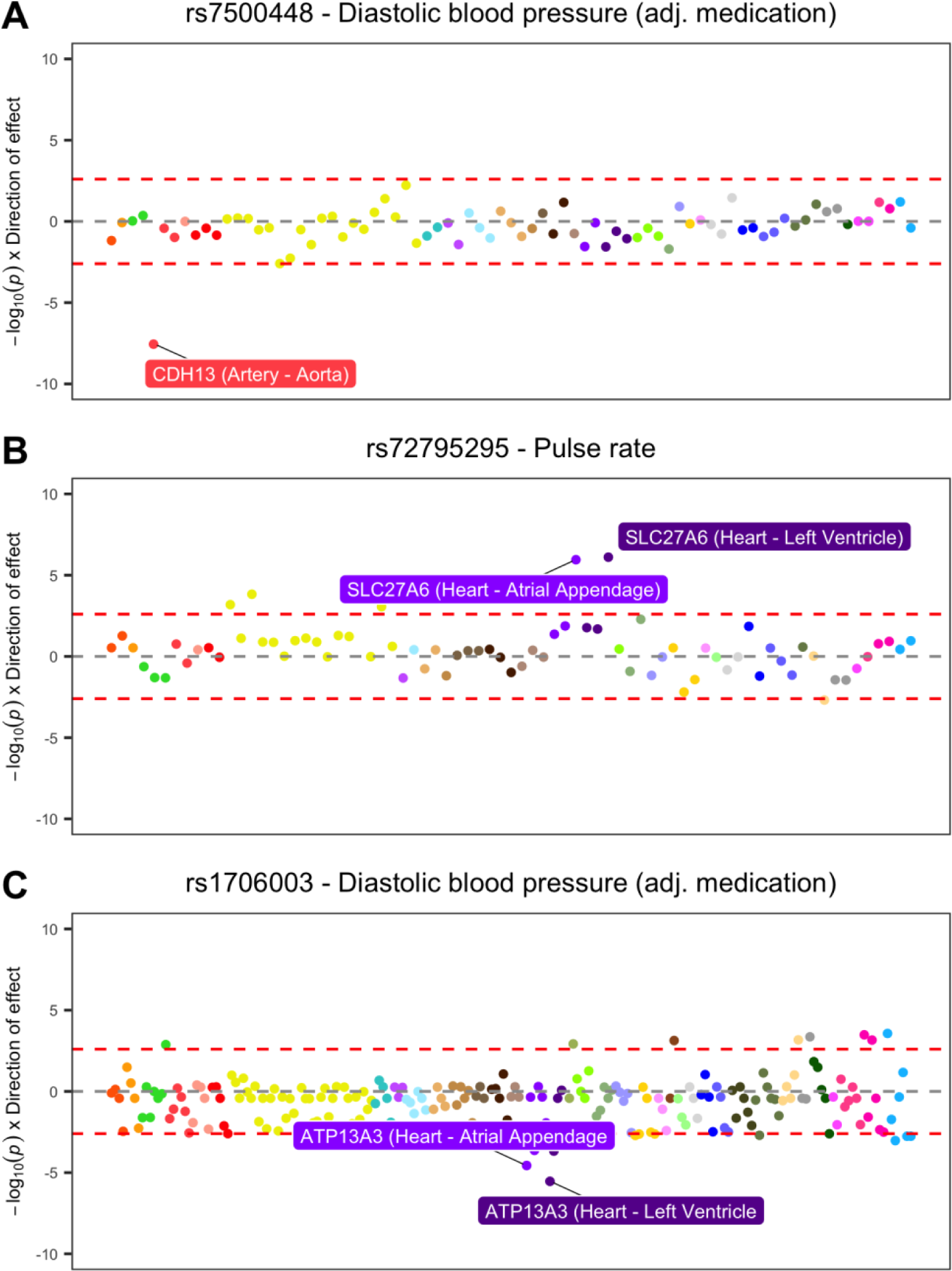
Miami plots between all genes whose expression is influenced by SNPs detected by genome-wide association studies (GWAS) of blood pressure traits. a) rs7500448 was strongly associated with diastolic blood pressure (DBP) based on *CDH13* expression derived from aorta tissue, b) rs72795295 was associated with pulse rate using *SLC27A6* heart-derived expression, c) rs1706003 was associated with DBP using *ATP13A3* expression data also derived from heart tissue.

This approach may also prove useful in identifying trait-associated variants yet to be discovered by GWAS. For instance, rs72795295 is most strongly associated with pulse rate out of the blood pressure related traits in our analysis, although this effect does not survive conventional GWAS multiple testing corrections (P=7.2 × 10^−08^). However, by integrating tissue-specific eQTL data, along with the reduced burden on multiple testing, our analysis provided evidence suggesting that this may be a novel trait-associated locus (Supplementary Table 20 and Figure 7b). Moreover, the strongest association in this evaluation was with *SLC27A6* expression derived from heart tissue (MR P=7.8 × 10^−07^), which again may help yield mechanistic insight into the causal pathway from genetic variant to phenotype.

Likewise, rs1706003 is a SNP associated with blood pressure which may be overlooked based on conventional GWAS corrections (P=1.1 × 10^−07^ with DBP). Integrating heart-derived eQTL data with findings from GWAS provided evidence which survived multiple comparisons in our analysis (MR P = 7.8 × 10^−07^) (Supplementary Table 21 and Figure 7c). Furthermore, although the nearest gene to rs1706003 is *TMEM44*, our results indicate that a gene located further downstream is more likely to be responsible for this effect (*ATP13A3*). Findings such as this support evidence that the nearest gene to a trait-associated SNP is not always the causative one^31^.

## Discussion

In this study we have undertaken a systematic phenome-wide association study to investigate the genetic effects of gene expression across different tissue types. In doing so, we have constructed a putative causal map of tissue-dependent associations across the human transcriptome. We have provided evidence that effects which influence gene expression across multiple tissue types are more likely to be associated with multiple traits. Our results also highlight the value of cross-tissue evaluations in terms of elucidating effects which depend upon the tissue analysed. We envisage that our findings will facilitate a greater understanding of tissue-specific regulatory mechanisms which are likely to have translational impact by informing drug target prioritization.

The tissues or cell types which a gene is expressed in is known to reflect the biological processes and functions it carries out^32^. For instance, in this study we demonstrated that the association between *TPO* and thyroid disease appears to be dependent on using expression data derived from thyroid tissue. This gene is responsible for generating thyroid peroxidase and thus plays an important role in regulating thyroid hormones^33^. As such this tissue-specific association reflects the role that this gene has in the thyroid gland. In contrast, the association between *RPS26* and thyroid disease was detected across all tissues evaluated. Moreover, the lack of heterogeneity detected in this cross-tissue evaluation suggests that the functional role of *RPS26* is set early in development. This gene encodes a ribosomal protein necessary for the production of 18S rRNA, a structural RNA which is a component of all eukaryotic cells^34^. Our results found that, along with thyroid disease, *RPS26* was linked with 47 other traits that survived multiple comparisons. The large number of identified effects therefore appears to reflect the function of *RPS26* which is likely crucial for many complex biological pathways. Broadly we also observed that variants which influence gene expression levels in multiple tissues are more likely to influence multiple complex traits. This suggests that genes expressed in many tissues are more likely to have widespread influence on downstream phenotypic consequences.

In our results we have demonstrated that phenome-wide evaluations of genes can help elucidate tissue-dependent associations. As an example of this, we show that *FBN2* is associated with various blood pressure traits when using expression data derived from heart tissue. However, when analysing *FBN2* expression using lung-derived data, these effects attenuated, whereas evidence of association with lung function and impedance were detected. This gene is responsible for encoding fibrillin 2 which is a glycoprotein responsible for elastin fibres found in connective tissue^35^. Elastin plays an important role in determining passive mechanical properties of the large arteries and lungs, which helps explain the associations detected in these separate tissues^36, 37^. *FBN2* is also associated with other traits and diseases, such as Marfan-like disorder^35^. A better understanding of pleiotropic effects due to regulatory mechanisms may also help to shed light on valid instruments in a conventional Mendelian randomization setting (i.e. between a modifiable environmental risk factor and disease outcome^8^). Specifically, an indication of number of genes’ expression that an instrument influences (and across how many diverse tissue types) would be valuable in a conventional MR setting.

Phenome-wide evaluations of our findings also have the potential to assist in drug target prioritisation. This supports emerging evidence concerning the benefit in using findings from genetic association studies to support therapeutic validation^38, 39^. Moreover, this is particularly crucial given the costs of drug development^40^, but also timely given that the highest number of novel drugs were approved in 2018^41^. As a proof of concept, we undertook a phenome-wide scan of *HMGCR* which is targeted by statins to reduce elevated cholesterol levels. We identified strong associations with cholesterol traits, but also findings which reflect known on-target effects of statins (namely changes in bodyweight and risk of diabetes^24^). So although GWAS datasets typically investigate disease incidence as opposed to disease progression or treatment, evaluations such as these may still be useful for therapeutic validation^23^.

Our results can also be used to flag on-target effects which are less well established in pharmacogenetics. For instance, our evaluation of *CYP19A1* suggested that inhibiting this target may result in lower bone mineral density. This finding supports a side-effect previously reported for the anti-cancer drug anastrozole which targets this gene^28^. The therapeutic benefit of statins on lower risk of coronary heart disease has been found to outweigh the adverse side effects on diabetes risk^42^. Uncovering potential side effects for other drug targets should motivate future endeavours to evaluate whether the benefits of therapeutic intervention outweigh the possible drawbacks. Similar evaluations may also help highlight potential drug repurposing and repositioning opportunities. We provide an example of this suggesting that targeting *ACHE* (originally targeted to treat cognitive decline in Alzheimer’s patients) may help lower blood pressure levels. There are likely many other potential associations from our analyses which may highlight potential drug repurposing/repositioning opportunities.

In the final series of analyses in our study, we propose that integrating tissue-specific eQTL data into GWAS analyses may help highlight genes responsible for association signals. Our approach therefore supports the notion of triangulation in epidemiology, whereby many lines of evidence are needed to support robust conclusions (i.e. colocalization of eQTL and GWAS effects)^43^. The examples we have showcased in this regard involve SNPs associated with blood pressure traits, where we prioritise *CDH13*, *SLC27A6* and *ATP13A3* as genes likely responsible for these effects. *CDH13* is a regulator of vascular wall remodelling and angiogenesis^44^, *SLC27A6* is responsible for a fatty acid transporter protein^45^ and *ATP13A3* has recently been implicated in pulmonary arterial hypertension susceptibility through rare loss of function analyses^46, 47^. However, although there are likely many instances where integrating tissue-specific eQTL data can help pinpoint genes responsible for GWAS associations, this may not always be possible due to the complexities of co-expression and widely expressed genes.

Endeavours which continue to generate increasingly large-scale tissue-specific molecular datasets will facilitate data mining opportunities across the human transcriptome^48^. Although the current sample sizes have meant that the analyses in this study have been restricted to using lead eQTL only, future efforts will benefit from leveraging multiple valid instruments within a Mendelian randomization framework. Nonetheless, techniques in genetic colocalization will likely continue to play an important role in discerning whether associations are detected due to shared causal variants. We also note that the inference of colocalization methods may be limited when evaluating associations at loci of dense linkage disequilibrium (such as the HLA region of the genome).

Furthermore, the approach used in our study (as with all alternatives to date) is unable to robustly rule out that findings may be influenced by molecular horizontal pleiotropy. This is the process whereby a genetic variant influences gene expression and a complex trait via two independent biological pathways. We also note that cross-tissue inference of our findings has the caveat of differing sample sizes in GTEx for different tissues. Lastly, when evaluating associations in our results it is important to remember that they are based on SNP effect sizes which are often relatively modest^49^, but potentially effective throughout the life course. Therefore, when evaluating our results for the purpose of drug validation it is worth noting that pharmaceutical targeting of a protein is likely to have a larger effect on protein levels, but over a shorter time period.

The results we have highlighted in our study are likely just the tip of the iceberg in terms of novel findings from our atlas that provide insight into the regulatory mechanisms underlying human complex traits. Although studies have used GTEx data to investigate tissue-specificity previously, their results are not easily accessible in a format that allow transcriptome-wide, phenome-wide or cross-tissue evaluations. Our web application should prove fruitful for users in this regard, facilitating in-depth evaluations of current findings or motivating innovative research hypotheses. Future endeavours which harness increasingly large-scale molecular datasets derived from different tissue types will enhance our capability to understand the determinants of complex disease.

## Methods

### Data resources

Tissue-specific eQTL data was obtained from the genotype-tissue expression (GTEx) project (v7) (https://gtexportal.org/home/). Only 48 of the 53 tissues available from GTEx v7 were analysed as each of the remaining 5 had fewer than 50 samples. We also obtained eQTL data derived from whole blood in 31,684 individuals made available by the eQTLGen consortium (http://www.eqtlgen.org). GWAS summary statistics were obtained from the Neale Lab analyses of UK Biobank data and consortia who have made their results publicly available (a full list can be found in Supplementary Table 2).

### Statistical analyses

We conducted analyses using the summary-data-based Mendelian randomization (SMR) method (v0.710). A reference panel of European individuals from the 1000 genomes project (phase 3) was used to compute LD estimation for all analyses^50^. As proposed previously^51^, only cis-eQTL were used as instrumental variables (based on < 1Mb of associated probe). This is to reduce the likelihood of associations attributed to horizontal pleiotropy to which trans-effects are more prone.

Consequently, only lead eQTLs for each gene were used as instrumental variables given that very few genes could be robustly instrumented with multiple independent SNPs in the GTEx dataset. This approach was also applied when analysing data from the eQTLGen consortium despite the larger sample sizes, for consistency when comparing associations between dataset. We defined eQTL based on a lenient p-value threshold of P < 1 × 10^−04^, maximizing the number of possible genes analysed across tissues but also allowing readers to filter out associations should they wish to apply a more stringent threshold.

An analysis of variance (ANOVA) model was applied to investigate the association between the number of traits and number of tissue types detected for all lead eQTL in our curated results (i.e. P < 5 × 10^−08^ that were also robust to HEIDI corrections). However, it is also possible that genomic properties (such linkage disequilibrium (LD) structure, proximity to nearest gene etc.) may influence the number of traits which multi-tissue eQTLs are associated with. Therefore, we adjusted our analysis for minor allele frequencies, linkage disequilibrium (LD) score and distance to gene expression probe for lead eQTL. Furthermore, associations detected using eQTLGen whole blood-derived data were removed from this analysis to reduce any bias which may be attributed to the large sample size of this dataset.

By default, our web application displays multiple testing comparisons based on Bonferroni correction for the number of tests undertaken in the search query. Subsequently HEIDI corrections are applied based on the number of associations which survived multiple testing in this look up.

All analyses were undertaken using R (version 3.5.1). The R package ‘shiny’ v1.1 was used to develop the web application. The R packages ‘manhattanly’ v0.2 and ‘highcharter’ v0.5 were used to generate interactive plots. Figures in this manuscript were generated using ‘ggplot2’ v2.2.1.

## Data availability

All results from the analyses undertaken in this study can be downloaded using our web application (http://mrcieu.mrsoftware.org/Tissue_MR_atlas/).

## Acknowledgements

We are extremely grateful to the GTEx, eQTLGen and GWAS consortia for making their summary statistics publicly available for the benefit of this study. This work was supported by the Integrative Epidemiology Unit which receives funding from the UK Medical Research Council and the University of Bristol (MC_UU_00011/1, MC_UU_00011/4 and MC_UU_00011/5). G.D.S, C.L.R and T.R.G conduct research at the NIHR Biomedical Research Centre at the University Hospitals Bristol NHS Foundation Trust and the University of Bristol. The views expressed in this publication are those of the author(s) and not necessarily those of the NHS, the National Institute for Health Research or the Department of Health. G.H is supported by the Wellcome Trust [208806/Z/17/Z]. T.G.R is a UKRI Innovation Research Fellow (MR/S003886/1).

## Materials and Correspondence

This publication is the work of the authors and T.G.R. will serve as guarantor for the contents of this paper.

